# Multiphase Organization Is a Second Phase Transition Within Multi-Component Biomolecular Condensates

**DOI:** 10.1101/2021.09.13.460104

**Authors:** Konstantinos Mazarakos, Huan-Xiang Zhou

## Abstract

We present a mean-field theory for the multiphase organization of multi-component biomolecular condensates and validate the theory by molecular dynamics simulations of model mixtures. A first phase transition results in the separation of the dense phase from the bulk phase. In a second phase transition, the components in the dense phase demix to localize in separate regions that attach to each other. The second phase transition occurs when the strength of cross-species attraction goes below the mean strength of the self-attraction of the individual species and reaches a critical value. At a given strength of cross-species attraction, both of the phase transitions can be observed by decreasing temperature, leading first to phase separation and then to demixing of the dense phase. The theory and simulations establish the disparity in strength between self and cross-species attraction as a main driver for the multiphase organization of multi-component biomolecular condensates.

**TOC GRAPHICS:** 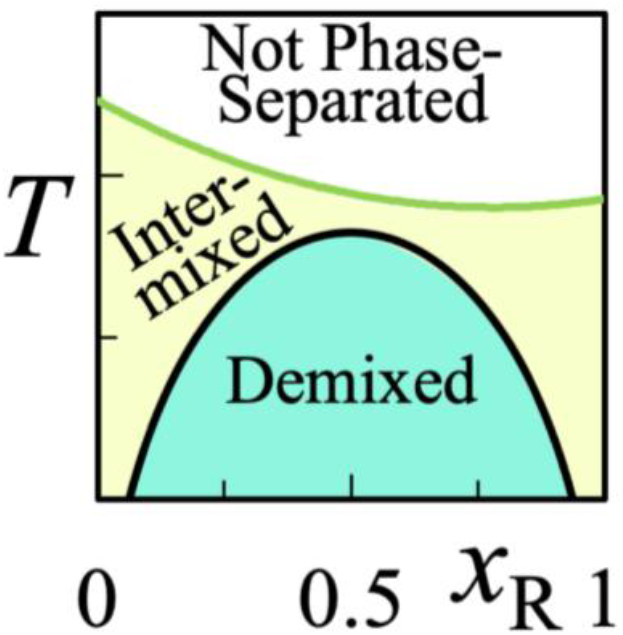

---

Biomolecular condensates, formed via liquid-liquid phase separation, mediate a myriad of cellular functions, including ribosome preassembly and mRNA sequestration under stress.^1-2^ Instead of a single homogeneous dense phase, multi-component condensates, both inside cells and reconstituted using purified components, show multiphase organization.^1, 3-11^ Multiphase coexistence has also been observed in coarse-grained molecular simulations.^8, 11-16^ Yet the theoretical underpinning is still unclear. Here we present a simple theoretical model to show that the multiphase organization of multi-component condensates is a second phase transition. Whereas the first phase transition that leads to the separation of condensates from the bulk phase is driven by overall attraction among the macromolecular components,^17^ the second phase transition, leading to multiphase organization within condensates, is driven by disparity in strength between self and cross-species attraction.

Our model systems have two components, D and R, which are either spherical particles or chains of such particles. The particles attract each other, with interaction energy −*ε*_*αβ*_ between species *α* and *β* (= D or R) at contact and 0 otherwise. The strengths, *ε*_DD_ and *ε*_RR_, of self-attraction of the two components, are different. Specifically, *ε*_DD_ > *ε*_RR_, so that the critical temperature for phase separation of pure D is higher than the counterpart of pure R. The phase separation of a mixture is driven by the self-attraction of D (for “driver”) but regulated by the self-attraction of R (for “regulator”) and the cross-species attraction between D and R.^14, 18-19^ We work in temperatures (*T*) below the critical temperature, *T*_c_, for the phase separation of the mixture, where the first phase transition has resulted in the coexistence of a dense phase and a bulk phase. Our interest is the dense phase, specifically its multiphase organization.

Let us first consider the case where the components are particles. In a mean-field treatment, the Helmholtz free energy, *F*, of the dense phase has an enthalpic contribution, due to pairwise interactions of the particles, and an entropic contribution, due to mixing of the species:

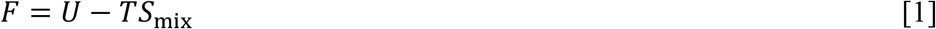

The enthalpic contribution can be obtained by enumerating the three kinds of contact pairs, DD, RR, and DR:

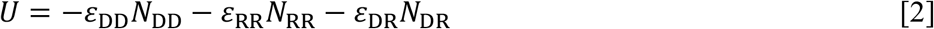

where *N*_*αβ*_ denotes the total number of contact pairs between species *α* and *β*. Let the total number of all contact pairs be *N*_p_:

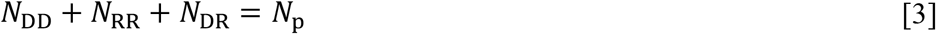

Note that *N*_p_ is proportional to the total number, *M*, of particles:

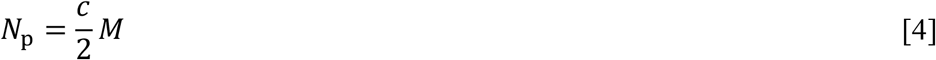

where *c* represents the average number of contact pairs formed by a particular particle.

Both the partition of *N*_p_ among the three kinds of contact pairs and the mixing entropy depend on the organization of the dense phase. Consider first the demixed state, where D and R form separate dense phases. We have

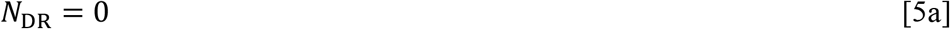

and

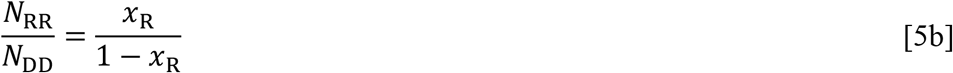

where *x*_R_ is the mole fraction of the regulator species. Solving eqs [3], [5a], and [5b], we find

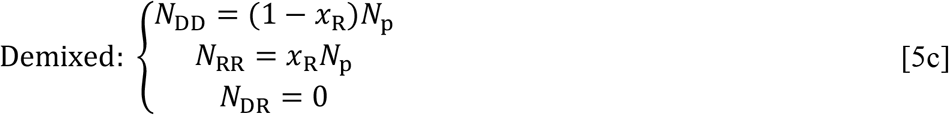

In the intermixed state, D and R form a single homogeneous dense phase. Around a given D particle, the chance of a contact partner being an R particle or another D particle is proportional to the mole fraction of the partner species. Thus

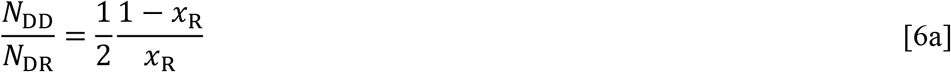

where the factor of one half accounts for the fact that DR pairs can be obtained in two ways: either D is the center and R is a contact partner, or R is the center and D is the contact partner. Likewise we have

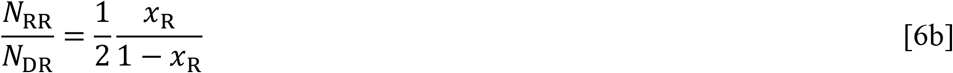

Solving eqs [3], [6a], and [6b], we find

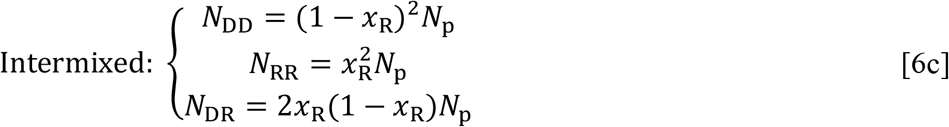

In the demixed state, by definition, the mixing entropy is 0,

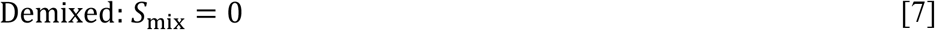

for any *x*_R_. Intermixing gives rise to a mixing entropy

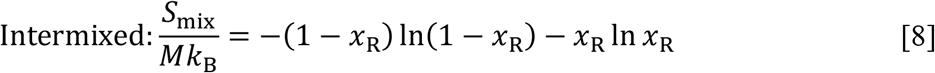

The change in *F* from the intermixed to the demixed state is finally found as^17^

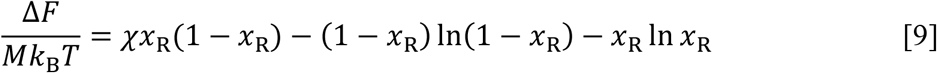

where

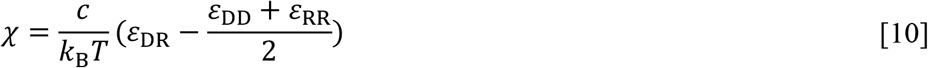

Decreasing *χ*, e.g., by weakening the cross-species attraction, favors the demixed state. When *χ* is decreased to a critical value,

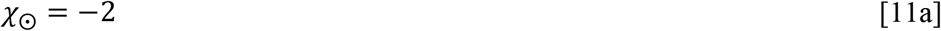

the demixed state becomes the thermodynamically stable state for the first time, at *x*_R_ = 0.5. If *ε*_DD_, *ε*_RR_, and *T* are fixed, the critical value for *ε*_DR_ is

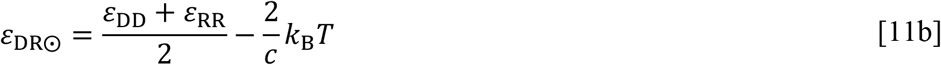

Note that *ε*_DR⨀_ is below the mean of *ε*_DD_ and *ε*_RR_, and the gap grows with increasing temperature. When *ε*_DR_ < *ε*_DR⨀_, the demixed state becomes stable (relative to the intermixed state) over a range of *x*_R_ around 0.5 (Fig. 1). For example, at *ε*_DR_ = *ε*_DR⨀_ − 0.1*k*_B_*T*/*c*, the stable *x*_R_ range of the demixed state is (0.315, 0.685). On the borders of this *x*_R_ range, the intermixed and demixed states are equally stable and hence they coexist. The relation between *ε*_DR_ and the two border *x*_R_ values is a binodal. If instead we fix *ε*_DR_ (below the mean of *ε*_DD_ and *ε*_RR_) and vary *T*, we obtain a critical temperature for demixing:

**Figure 1.**
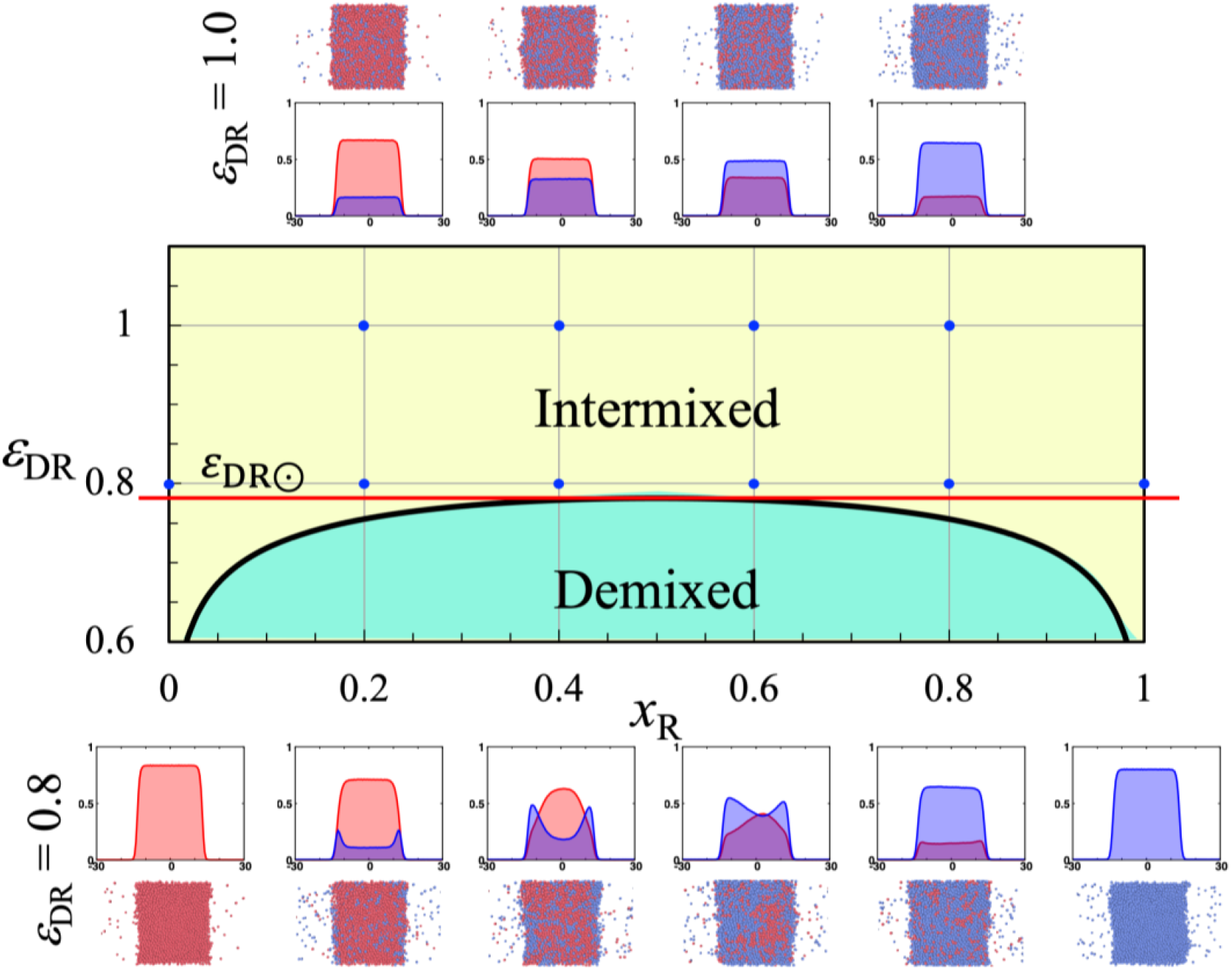
The *ε*_DR_ vs. *x*_R_ binodal of particle mixtures predicted by the mean-field theory. The binodal separates the intermixed zone (yellow) from the demixed zone (cyan). The critical value for *ε*_DR_ is shown by a red horizontal line. Density profiles of the two species (D: red; R: blue) and representative snapshots of particle mixtures obtained in molecular dynamics simulations are shown for the indicated *ε*_DR_ and *x*_R_ values.

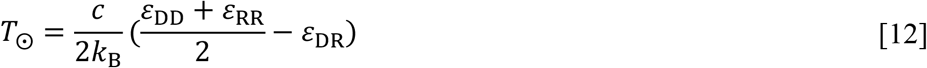

When *T* < *T*_⨀_, there is a range of *x*_R_ values, centered at *x*_R_ = 0.5, in which the demixed state is stable. The relation between *T* and the two border *x*_R_ values is another type of binodal.

The above free energy calculation can be easily extended to polymer blends, where each polymer is a chain of particles. For a binary blend, with both component the same chain length *L* (representing the number of particles per chain), the free energy of the dense phase is^20-21^

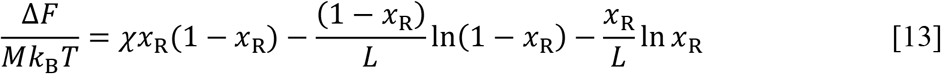

where the mixing entropy is reduced due to chain connectivity. The critical *χ* value for demixing is now

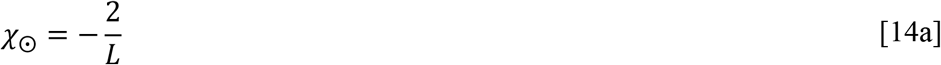

Chain connectivity thus reduces the magnitude of *χ*_⨀_. The corresponding critical value for *ε*_DR_ is

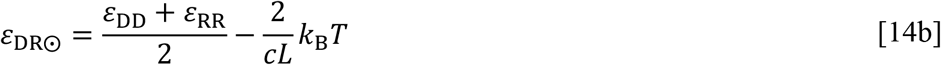

A comparison between eqs [11b] and [14b] shows that the gap between *ε*_DR⨀_ and the mean of *ε*_DD_ and *ε*_RR_ is narrower for chain mixtures than for particle mixtures. The critical temperature for chain mixtures is

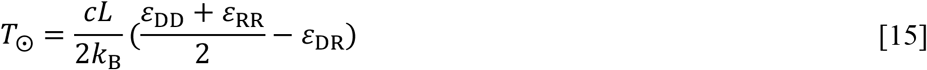

which is higher than the counterpart (eq [12]) for particle mixtures. The higher critical values of *ε*_DR_ and temperature mean that demixing occurs more easily in chain mixtures than in particle mixtures.

This mean-field theory makes a number of important predictions on the organization of the dense phase of mixtures formed by phase separation. First, when the strength (*ε*_DR_) of the cross-species attraction decreases below the mean strength of the self-attraction of the individual species and reaches a critical value *ε*_DR⨀_, a second phase transition occurs within the dense phase, resulting in demixing of the species. Second, the second phase transition occurs at a higher *ε*_DR_ for chain mixtures than for particle mixtures. Third, when *ε*_DR_ is below the critical value *ε*_DR⨀_, there is a range of mixing ratios, centered at equimolar mixing (i.e., *x*_R_ = 0.5), where demixing occurs. This means that demixing most readily occurs when the species are mixed at equimolarity. Fourth, when *ε*_DR_ is below the mean of *ε*_DD_ and *ε*_RR_, there is a critical temperature *T*_⨀_. Demixing occurs only when *T* is below *T*_⨀_. All these predictions are consistent with our initial results of demixing from molecular dynamics simulations of binary mixtures of Lennard-Jones particles or chains of such particles (*L* =10).^14^ For these Lennard-Jones systems, we used *ε*_*αβ*_to denote the well-depth for the interaction potential energy between particles of species *α* and *β* (= D or R), and fixed *ε*_DD_ and *ε*_RR_ at 1 and 0.9, respectively. Complete intermixing between D and R was observed at *ε*_DR_ = 1.2 and 1.0, but demixing was observed at *ε*_DR_ = 0.8 for chain mixtures at lower temperatures. We thus suspected that demixing was driven by the disparity between cross-species interaction strength *ε*_DR_ and the self-interaction strengths *ε*_DD_ and *ε*_RR_. This suspicion is now justified by the mean-field theory. We now reexamine the simulation results on the demixing of the Lennard-Jones systems in light of the mean-field theory. Demixing slows down the equilibration process and also can produce multiple metastable states. We thus extended the simulations here from 10 million to 100 million steps (ref ^14^; Fig. S1) and carried out five replicate runs when there were signs of demixing.

In Fig. 1, we display the density profiles and representative snapshots of the particle mixtures at *k*_B_*T* = 0.65, along with the *ε*_DR_ vs. *x*_R_ binodal predicted by the mean-field theory. As already noted, at *ε*_DR_ ≥ 1.0, only the intermixed state was observed, as indicated by uniform densities of the components inside the dense phase. At *ε*_DR_ = 0.8, there was local enrichment of the regulator (i.e., lower-*T*_c_) species at the interface between the dense and bulk phases, which we attribute at least partly to the finite size of the simulation systems and does not necessarily indicate the demixed state per se. With *c* = 7.7, eq [11b] predicts *ε*_DR⨀_ = 0.78, which is consistent with the absence of demixing in the simulations at *ε*_DR_ = 0.8. The mean-field theory further predicts demixing at *ε*_DR_ = 0.6 for all *x*_R_ values except those near 0 or 1, which are in agreement with simulations (not shown).

We display the corresponding results for chain mixtures at *k*_B_*T* = 1.7 in Fig. 2. Again, only intermixing was observed at *ε*_DR_ = 1.0, but now there was clear indication of demixing at *ε*_DR_ = 0.8 for 0.2 ≤ *x*_R_ ≤ 0.8. These observations are consistent with the mean-field theory, which predicts *ε*_DR⨀_ = 0.84 (eq [14b] with *cL* = 31) and demixing for 0.105 ≤ *x*_R_ ≤ 0.895 at *ε*_DR_ = 0.8. At *x*_R_ = 0.2 and 0.4, the dense phase in the simulations consisted of a D-rich region at the center, bordered by two R-rich regions on the two sides (“R-D-R” configuration; bottom rows in Figs. 2 and S2A,B). At *x*_R_ = 0.6, the dense phase had the R-D-R configuration in two of the five replicate runs but a “D-R” configuration in the other three runs, where a D-rich region and an R-rich region attach to each other (bottom row in Figs. 2 and S2C). At *x*_R_ = 0.8, the D chains in the dense phase coalesce into a cylinder, located near the interface with the bulk phase (bottom row in Figs. 2 and S2D).

**Figure 2.**
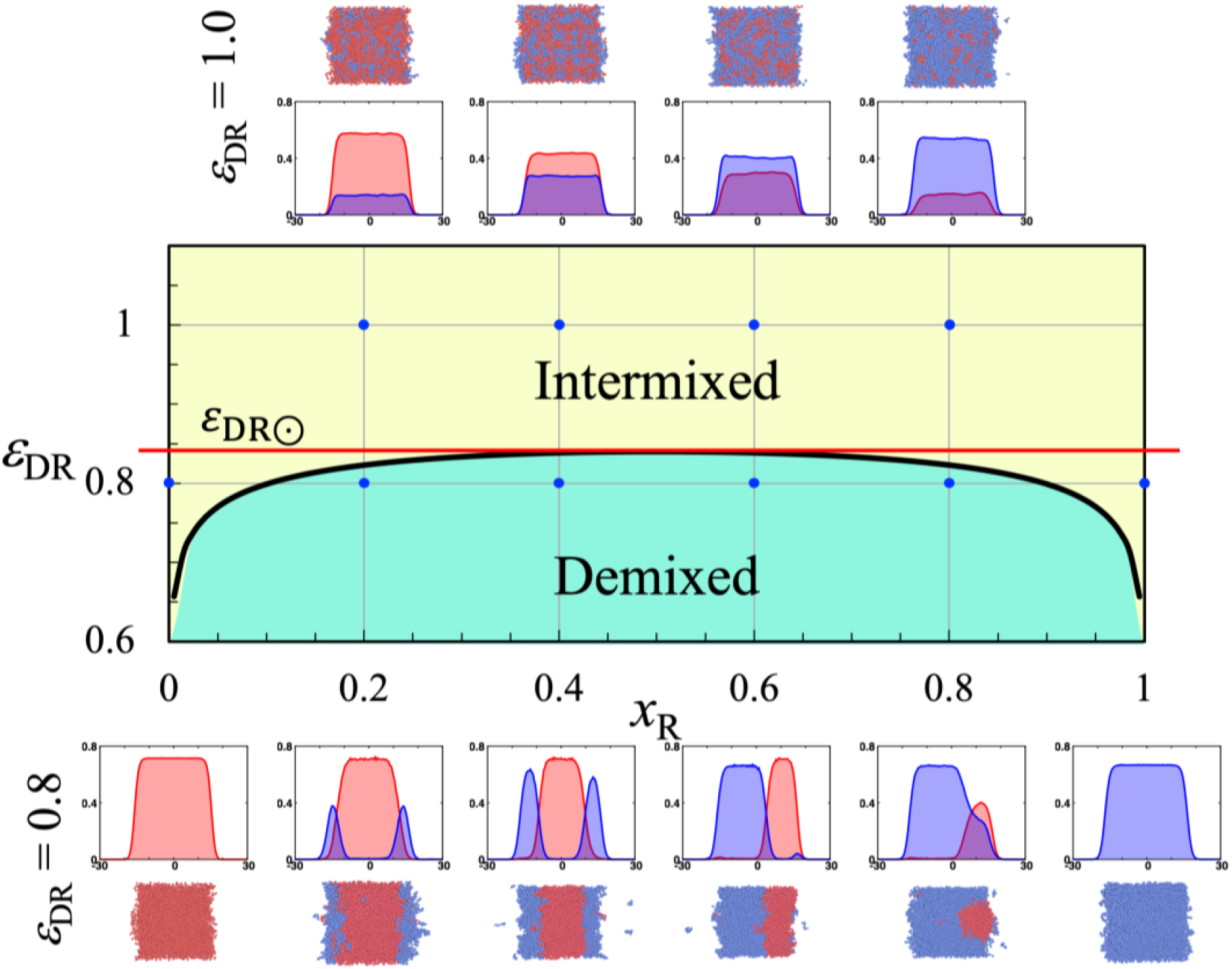
The *ε*_DR_ vs. *x*_R_ binodal of chain mixtures predicted by the mean-field theory. The binodal separates the intermixed zone (yellow) from the demixed zone (cyan). The critical value for *ε*_DR_ is shown by a red horizontal line. Density profiles of the two species (D: red; R: blue) and representative snapshots of chain mixtures obtained in molecular dynamics simulations are shown for the indicated *ε*_DR_ and *x*_R_ values.

Next we consider the effects of temperature on the behaviors of the dense phase of chain mixtures at *ε*_DR_ = 0.8. Figure 3 displays the *T* vs. *x*_R_ binodal predicted by the mean-field theory, with a critical temperature of *k*_B_*T*_⨀_ = 2.325 for demixing. It should be noted that there is also a critical temperature, *T*_c_, for phase separation (i.e., the first phase transition). The *k*_B_*T*_c_ values have a parabolic dependence on *x*_R_, with a minimum of 2.403 at *x*_R_ = 0.78 (green curve in Fig. 3).^14^ The entire *T* vs. *x*_R_ plane can be divided into three distinct zones. Above the *T*_c_ curve, there is no phase separation; between the *T*_c_ curve and the *T* vs. *x*_R_ binodal, the dense phase is a homogenous mixture of the components; below the *T* vs. *x*_R_ binodal, the dense phase is in a demixed state. As already noted when presenting Fig. 2, when *k*_B_*T* = 1.7, the chain mixtures with *x*_R_ = 0.2, 0.4, 0.6, and 0.8 all fell into the demixed zone. In contrast, when *k*_B_*T* = 2.3, these mixtures all fell into the intermixed zone (top row in Fig. 3). At this temperature, the two species were well-mixed in the dense phase, although for *x*_R_ = 0.2 and 0.4 there was local enrichment of R chains at the interface between the dense and bulk phases, which we again attribute to the finite size of the simulation systems.

**Figure 3.**
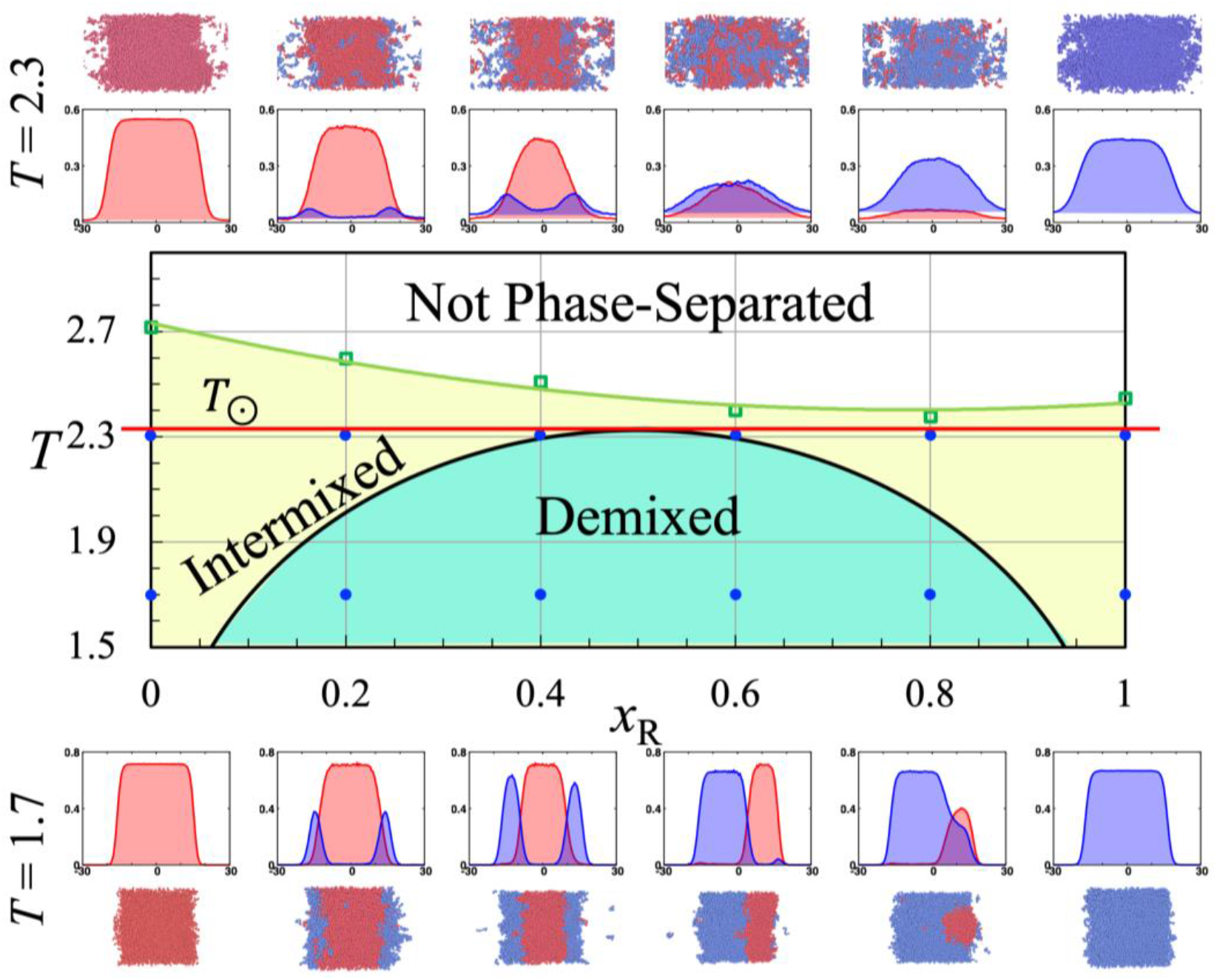
The *T* vs. *x*_R_ binodal of chain mixtures predicted by the mean-field theory. The binodal (black curve) separates the intermixed zone (yellow) from the demixed zone (cyan). The *T*_c_ curve (green) further separates the intermixed zone from the non-phase-separated zone (white). A red horizontal line is drawn at *T* = *T*_⨀_ (in units of 1/*k*_B_). Density profiles of the two species (D: red; R: blue) and representative snapshots of chain mixtures obtained in molecular dynamics simulations are shown for the indicated *T* and *x*_R_ values.

Lastly let us look at the transition from the demixed state to the intermixed state at a fixed *x*_R_ as *k*_B_*T* increased from 1.7 to 2.3 (Figs. 4 and S2). For *x*_R_ = 0.2 and 0.4, with increasing temperature, more and more R chains migrated from the outer R-rich regions to the central D-rich region, leading to the merge of the R-rich regions, while D chains migrated in the reverse direction. The same scenario applied to the R-D-R configuration at *x*_R_ = 0.6 when *k*_B_*T* changed from 1.7 to 2.3. For the D-R configuration at *x*_R_ = 0.6 and *k*_B_*T* = 1.7, with increasing temperature, D-chains and R-chains migrated toward each other, ultimately leading to the complete mixing of the two species. For *x*_R_ = 0.8, as *k*_B_*T* changed from 1.7 to 2.0, more and more D-chains in the cylinder migrated into the R-rich region, while R-chains spread into a wider region. When *k*_B_*T* reached 2.1, the two species were already well-mixed.

**Figure 4.**
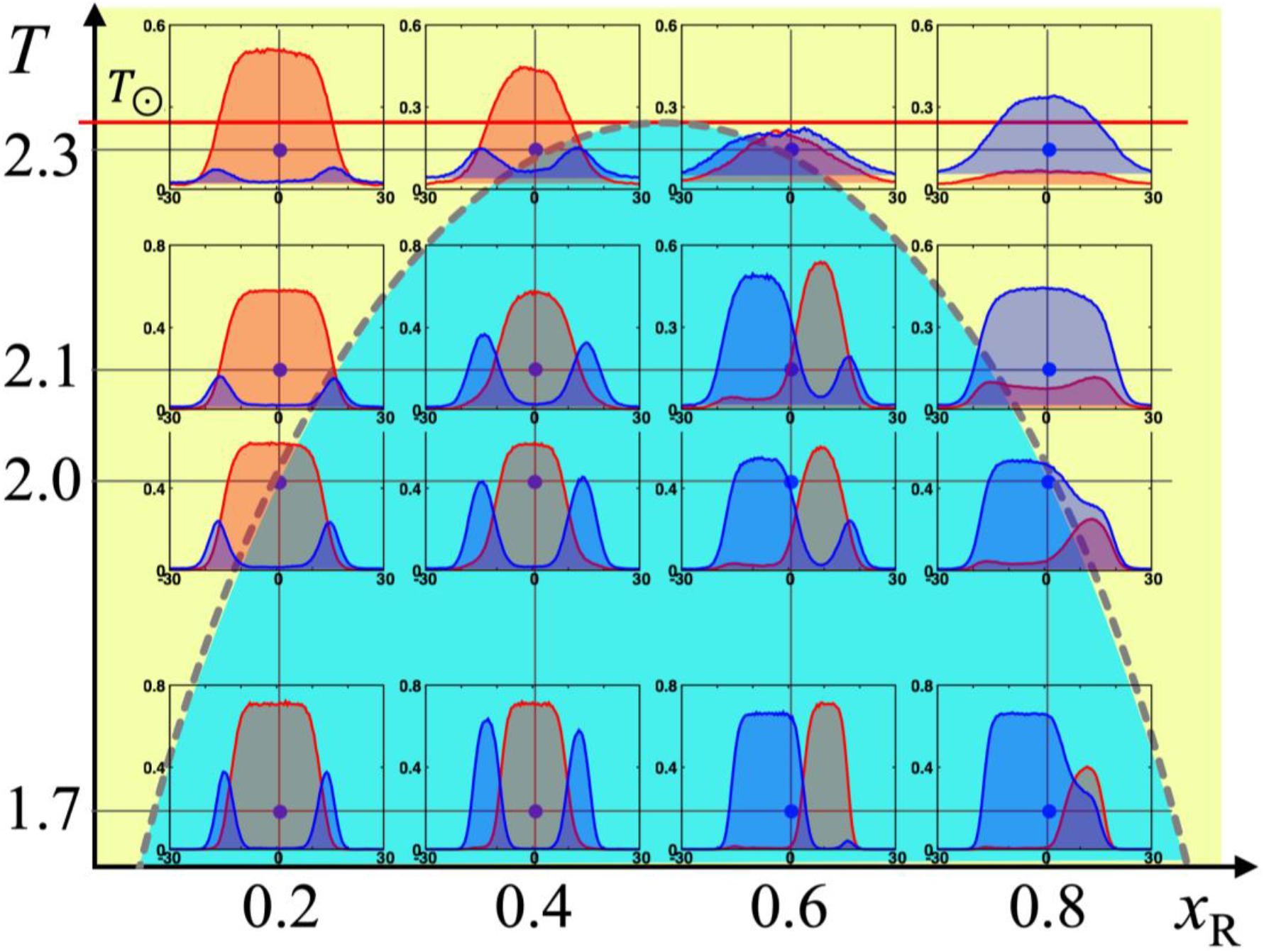
Detailed view into an area spanning *k*_B_*T* from 1.7 to 2.3 and *x*_R_ from 0.2 to 0.8, showing the transition from the demixed at *k*_B_*T* = 1.7 to the intermixed state at *k*_B_*T* = 2.3. Density profiles of the two species (D: red; R: blue) and representative snapshots of chain mixtures obtained in molecular dynamics simulations are shown for various combinations of *T* and *x*_R_ values.

In conclusion, we have presented a mean-field theory that demonstrates that the multiphase reorganization in the dense phase of mixtures is a second phase transition, after the first phase transition resulting in separation of the dense phase from the bulk phase. Molecular dynamics simulations of model mixtures have provided crucial test of the theory, including the existence of a critical value for the strength of cross-species attraction and a critical temperature for demixing. At a given strength of cross-species attraction, both of the phase transitions can be observed by decreasing temperature, leading first to phase separation and then to demixing of the dense phase. The theory and simulations establish the disparity in strength between self and cross-species attraction as a main driver for multiphase organization, and provide a much needed conceptual framework for understanding the complex phase behaviors of bimolecular condensates.

## ASSOCIATED CONTENT

### Supporting Information

The following file is available free of charge.

Two supporting figures: illustrating the procedure of the molecular dynamics simulations; showing an expanded version of Fig. 4, with results from five replications for each (*T, x*_R_) combination (PDF)

## AUTHOR INFORMATION

### Notes

The authors declare no competing financial interests.

## ACKNOWLEDGMENT

This work was supported by National Institutes of Health Grant GM118091.

## Notes

### Competing Interest Statement

The authors have declared no competing interest.

