## Supporting Figures for "Multiphase Organization Is a Second Phase Transition Within Multi-Component Biomolecular Condensates"

AUTHOR INFORMATION

**Corresponding Author**

*

*Supporting Information*

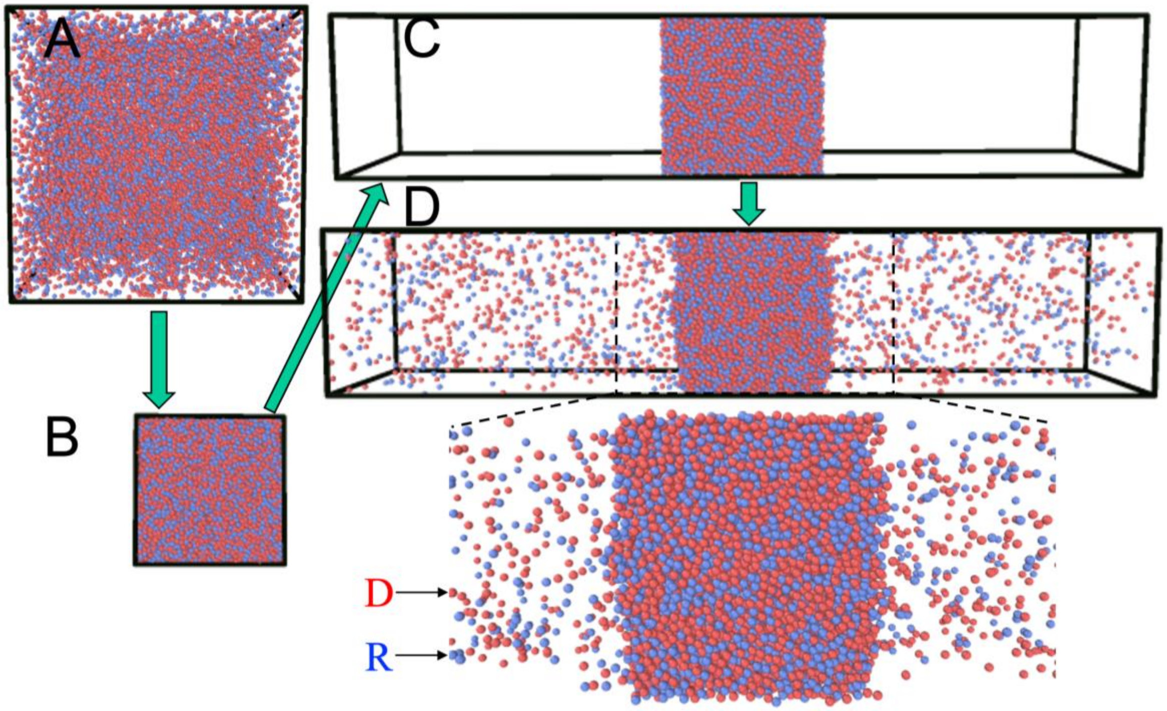


**Figure S1.** Illustration of the simulation procedure. (A) System in an initial cubic box. (B) System after two-fold compression in each direction. (C) Enlarged box in *z* direction. (D) Two-phase equilibrium. A zoomed view of the dense phase and the neighboring regions in the bulk phase is shown at the bottom. This figure has appeared in ref. 11.

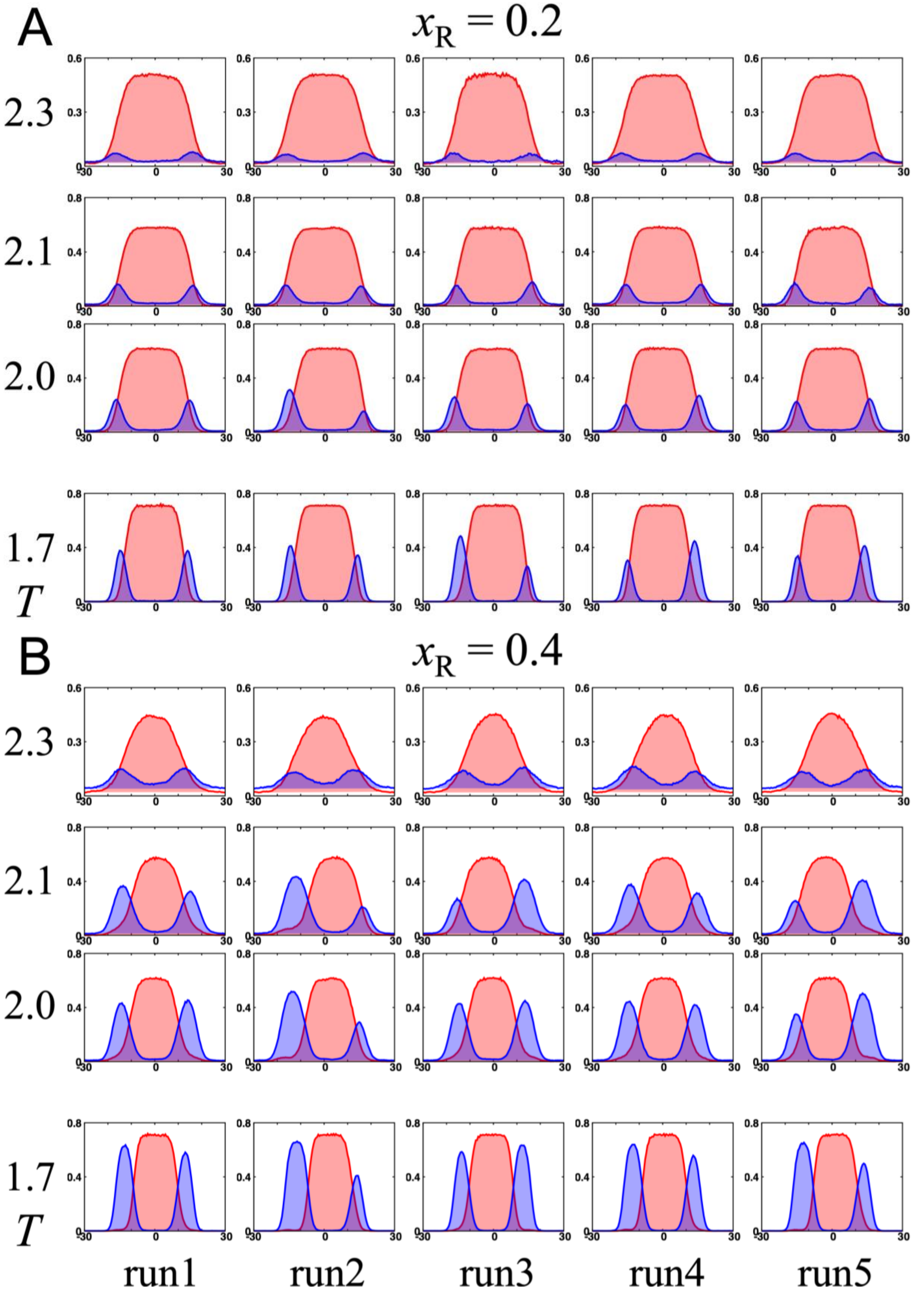
A
$x_R = 0.2$
2.3
2.1
2.0
1.7
$T$
B
$x_R = 0.4$
2.3
2.1
2.0
1.7
$T$
run1
run2
run3
run4
run5

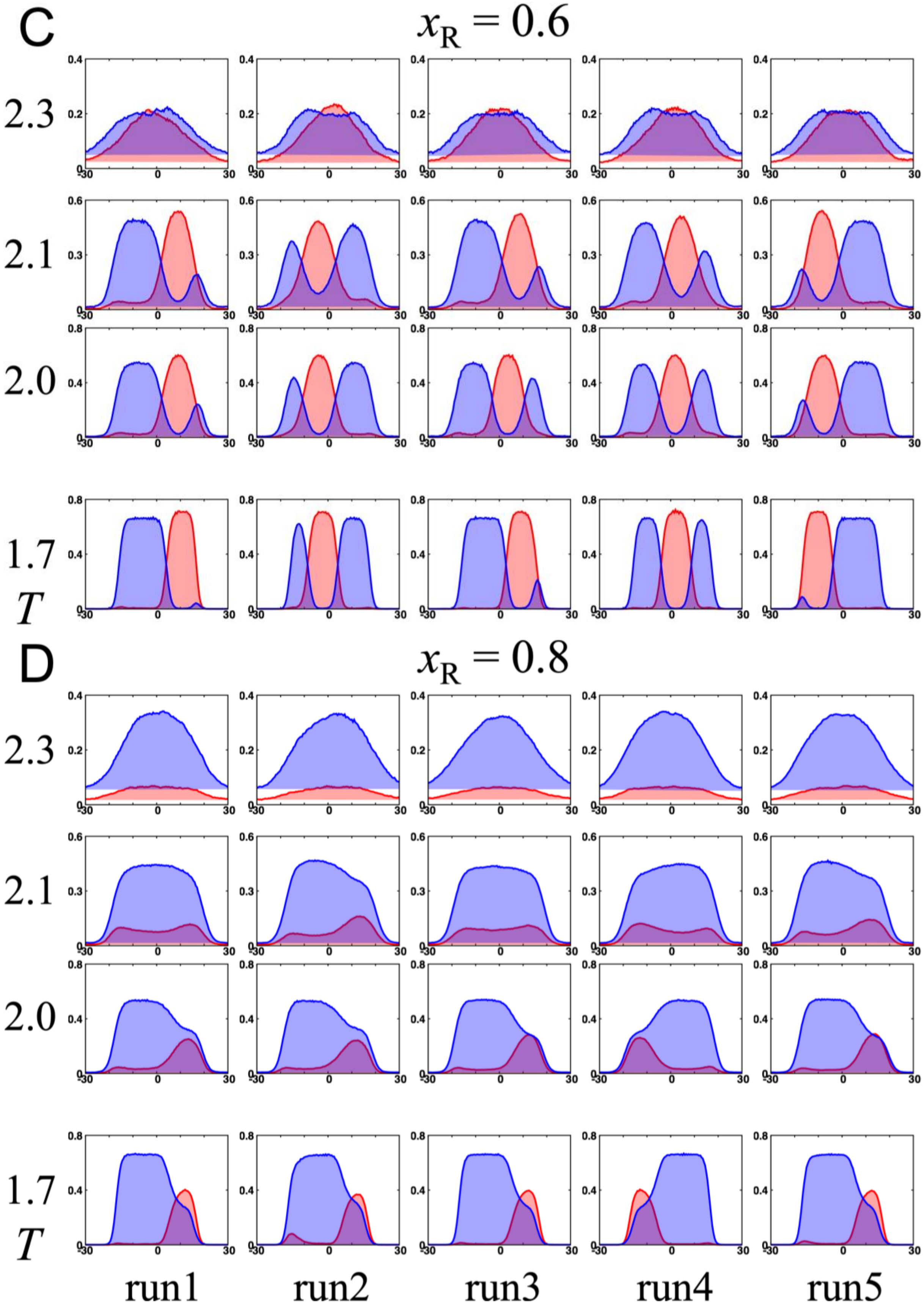
C
$x_R = 0.6$
2.3
2.1
2.0
1.7
T
D
$x_R = 0.8$
2.3
2.1
2.0
1.7
T
run1
run2
run3
run4
run5

**Figure S2.** Density profiles of the two species (D: red; R: blue) in molecular dynamics simulations of chain mixtures, from five replicate runs at $k_B T$ = 1.7, 2.0, 2.1, and 2.3 and a range of $x_R$ values. (A-D) $x_R$ = 0.2, 0.4, 0.6, and 0.8. Results for run1 are also shown in Fig. 4.
